# Temporal dynamics of early somatosensory processing in goal-directed actions

**DOI:** 10.64898/2026.01.16.699864

**Authors:** Elena Fuehrer, Dimitris Voudouris, Lisa K. Maurer, Katja Fiehler

**Author notes:** Correspondence: Katja Fiehler.

## Abstract

Tactile sensitivity is temporally modulated on a moving limb. Yet the neural mechanisms underlying this modulation remain unclear. Using electroencephalography, we examined the temporal dynamics of early somatosensory processing during goal-directed reaching. Participants reached for their unseen left hand while brief vibrations were applied to the moving right index finger in different movement phases. Psychophysical results revealed overall tactile suppression during reaching, with sensitivity transiently improving around maximum speed relative to the late movement phase. Consistent with the psychophysical data, the short-latency P45 component, originating in primary somatosensory cortex, was suppressed during reaching but recovered around maximum speed. Reduced P45 amplitudes were associated with stronger tactile suppression measured behaviorally, linking early cortical activity to perceptual changes. This temporally specific modulation of tactile perception, associated with early cortical processing, suggests that the somatosensory system dynamically adapts its sensitivity to facilitate sensorimotor control in goal-directed actions.

**Significance statement:** The perception of touch on a moving hand varies across different movement phases, depending on the importance of sensory feedback for guiding the hand. Our results show that these changes in perception are linked to modulations in early brain activity, indicating that the central nervous system adjusts touch processing in a time-sensitive manner at early processing stages to facilitate control of goal-directed actions.

## Introduction

Goal-directed movements depend on somatosensory signals. For example, tactile sensations directly arise from physical interactions involving the moving limb, and proprioceptive and tactile feedback inform about limb position and movement dynamics. Some of these somatosensory signals provide essential information for sensorimotor control (Goulding et al., 2025), whereas others may be irrelevant. Because feedback-based mechanisms alone are inherently noisy (Faisal et al., 2008) and temporally delayed (Miall & Wolpert, 1996), the sensorimotor system likely integrates sensory feedback with predictive feedforward models using efference copies of motor commands to estimate future sensory states of the moving limb (Desmurget & Grafton, 2000).

A well-documented perceptual phenomenon related to sensorimotor control is a reduction in tactile sensitivity on the moving limb, known as tactile suppression (Chapman et al., 1987). This phenomenon has been demonstrated by behavioral studies across a wide range of actions and is complemented by electroencephalographic (EEG) findings, showing reduced amplitudes of somatosensory evoked potentials (SEPs) during movement, particularly at short-latency components like the N20 and P45 (Lei et al., 2018; Rossini et al., 1999; Rushton et al., 1981), associated with early-stage processing in primary somatosensory cortex (S1) (Allison et al., 1992).

Crucially, tactile suppression is not an all-or-nothing phenomenon: When sensory input from the moving limb is relevant to successfully perform the movement, tactile sensitivity is preserved to some extent, although partial suppression may persist. For example, tactile suppression is less pronounced on the index finger when grasping an object with a precision grip, as opposed to other parts of the hand and arm (Colino et al., 2014; Manzone et al., 2018), or when grasping a slippery versus a high-friction object without seeing it (Voudouris & Fiehler, 2022). Tactile perception also recovers on the lower leg shortly before dealing with perturbations during a balancing task (Wachsmann et al., 2025b). Likewise, tactile suppression is evident during targeted reaching movements without visual feedback, which rely exclusively on somatosensory inputs (Voudouris & Fiehler, 2021). Importantly, tactile suppression during such reaching actions follows distinct temporal dynamics, as it is transiently released around the time of maximum speed (Voudouris & Fiehler, 2021), i.e., the onset of the deceleration phase when sensory guidance of the movement becomes particularly important (Elliott et al., 2001). Together, these findings demonstrate a flexible modulation of tactile suppression according to the functional relevance of somatosensory feedback. However, because these observations are based on behavioral measures only, it remains unclear whether changes in tactile perception reflect corresponding neural modulations in early somatosensory processing.

Electrophysiological studies provide some evidence for relevance-based modulation of SEPs from the moving limb: short-latency SEP amplitudes increase during exploratory hand movements compared to rest (Knecht et al., 1993), and during lower limb movements in a balancing task when an upcoming perturbation challenges posture (McIlroy et al., 2003). In a magnetoencephalographic study, the somatosensory evoked field component M38 was enhanced during grasping movements relative to rest and further increased during a more complex ball rotation task (Wasaka et al., 2017). While these electrophysiological studies did not measure psychophysical aspects of tactile perception, they suggest that facilitation of early somatosensory processing may underlie the modulation of tactile suppression during movement as sensory relevance changes.

To jointly examine the behavioral and neural correlates of this dynamic modulation, we combined psychophysics with EEG recordings of SEPs during a reaching task previously shown to modulate tactile perception (Voudouris & Fiehler, 2021). First, we asked whether tactile sensitivity is modulated across different movement phases; second, whether early cortical processing indexed by the P45 component shows a corresponding temporal modulation; and third, whether temporal changes in tactile sensitivity are linked to changes in cortical processing. Based on prior evidence (Rossini et al., 1999; Rushton et al., 1981), we predicted that movement-related tactile suppression would correspond to reductions in the early P45 component and that the time course of P45 amplitudes would mirror the temporal dynamics of suppression across movement phases.

## Materials and Methods

### Experimental Design

Vibrations (20 ms, 280 Hz) were delivered using a vibrotactile stimulation device (PiezoTac tactor; Engineering Acoustics Inc., Casselberry, FL, USA). The tactor comprised a skin contactor (6.4 mm diameter) mounted on a shielded cantilever. Cantilever vibrations produced sinusoidal oscillations perpendicular to the skin. One contactor was placed on the palmar side of the proximal phalanx of the right index finger, leaving the fingertip uncovered. A second contactor was positioned on the sternum, approximately 3 cm below the clavicle. The contactors were fixed to the participant’s skin using medical tape. The experiment was controlled by custom MATLAB (The MathWorks Inc., Natick, MA, USA) software using the Psychtoolbox extension (Brainard, 1997). The position of an infrared marker on the nail of the right index finger was recorded at 250 Hz with an Optotrak Certus motion tracking system (Northern Digital Inc., Waterloo, ON, Canada).

### Procedure

Participants placed both hands on a table in front of them. The left hand rested palm-down with the thumb and index finger on specific markers positioned 10 cm to the left of the body midline, to ensure consistent placement. The right hand rested on a keypad located 18 cm to the right of the midline and 20 cm in front of the participant. Throughout the experiment, participants’ view of their hands was occluded using a tilted cardboard screen, which allowed them to see the top of the computer monitor for instructions. At the beginning of each trial, participants held down a key with their right index finger to initiate the trial. Text on the computer screen then indicated the target finger on the left hand (thumb or index), and participants had 2500 ms to initiate the movement.

Upon releasing the key, participants performed a reaching movement with the right hand to touch the nail of the specified target finger. A standard vibration (peak-to-peak displacement: 119 µm) was delivered to the moving right index finger at one of five possible latencies relative to movement onset (50, 150, 250, 350, or 450 ms), which was determined by the key release. After reaching the target, participants rested their right hand next to the left hand on the table. A comparison vibration was delivered to the sternum 3000 ms after the standard vibration, well after movement completion, while both hands were still. The comparison stimulus had one of seven possible intensities (peak-to-peak displacement: 47, 71, 95, 119, 143, 167, or 192 µm). Participants indicated which of the two vibrations felt more intense by pressing foot pedals: the left pedal for the first vibration (on the hand) and the right pedal for the second vibration (on the sternum; see Fig. 1 for the trial sequence). Each stimulation time was presented 70 times for a total of 350 trials. Additionally, 70 trials (16.67% of all trials) were included to record movement-related EEG without vibrotactile stimulation, matching the number of trials for each stimulation time. In these omission trials, participants were informed that no vibration was given 3000 ms after movement onset and no tactile discrimination response was required. Participants completed 10 blocks of 42 trials each. Within each block, all combinations of stimulation time and comparison intensity were presented once, along with seven omission trials in random order. Trials were repeated at the end of the block if participants initiated the movement too early, less than 180 ms after target presentation, suggesting the cue had not been processed, or if they failed to initiate movement within 2500 ms after the cue. Self-paced breaks were provided between blocks. Before starting the experiment, participants familiarized themselves with the reaching movement and the discrimination task in 20–40 practice trials. We also assessed tactile sensitivity at rest by asking participants to perform the discrimination task without movement, while both hands remained on the table in the same position adopted after the reaching movements. This enabled us to assess relative changes between rest and movement phases and determine the effect of movement on tactile processing while accounting for idiosyncrasies in tactile sensitivity. The standard vibration was presented to the right index finger at one of five delays (50, 150, 250, 350, or 450 ms) after a fixed interval of 700 ms from trial onset, replicating the relative timing used during movement. A comparison vibration followed on the sternum 3000 ms later. Participants judged vibration intensity using foot pedals. Each of the seven comparison intensities was presented twice per stimulation time (70 trials total). Trials ended only after participants had responded and at least 6000 ms had elapsed from trial onset, ensuring a trial duration consistent with the reaching block. Trials started automatically after a foot pedal was pressed. The rest condition was divided into two blocks, separated by a short break, and both blocks were administered either before or after the movement blocks, counterbalanced across participants.

**Fig. 1:**
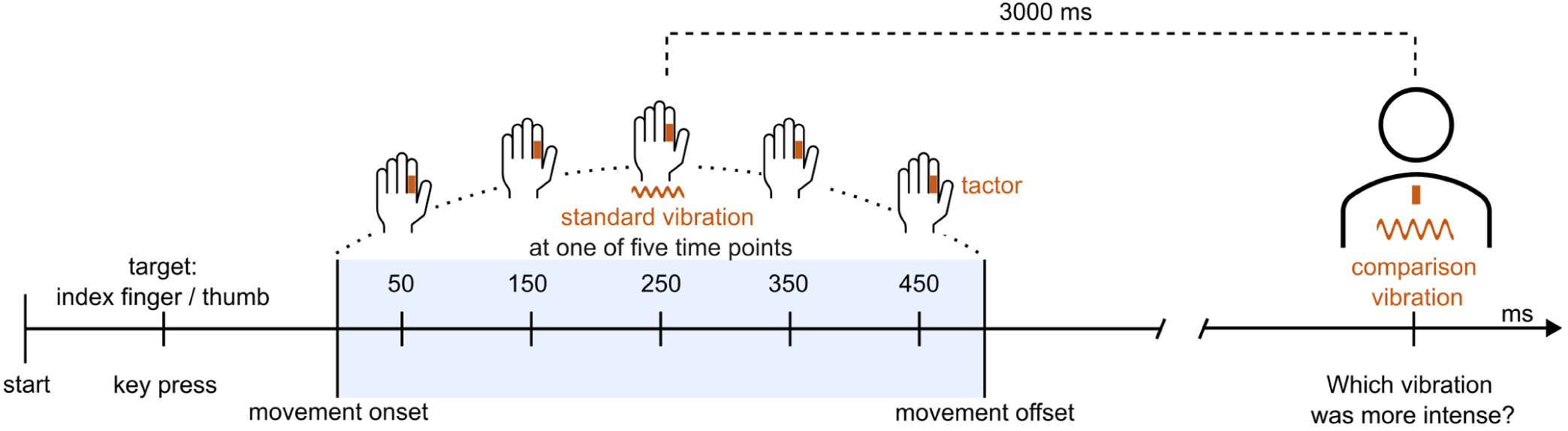
Trial Sequence. Participants initiated each trial by pressing and holding a key with the right index finger. Text on the monitor indicated the target finger of the left hand. Participants released the key and reached toward the target finger of the left hand. A standard vibration of fixed amplitude was delivered to the right index finger at one of five time points during the movement (250 ms shown here). After an ISI of 3000 ms, a comparison vibration of varying amplitude was presented on the sternum. Participants judged which vibration felt stronger using foot pedals. Participants also performed the same psychophysical task with both hands at rest.

### Participants

Thirty-eight participants (self-reported gender: 22 female, 16 male; mean age: 25.7 years, SD = 4.1) provided written informed consent and received either course credit or 10 €/hour as compensation. The study was approved by the ethics committee of the Justus Liebig University Giessen and conducted in accordance with the Declaration of Helsinki (World Medical Association, 2013), with the exception of pre-registration. All participants were right-handed, as assessed by the Edinburgh Handedness Inventory(Oldfield, 1971) (M = 90.8, range = 28.6–100). Two participants were excluded for slow movements, leaving too few stimulus presentations in the late post-max speed phase to fit psychometric functions (see Kinematic Analysis); one participant was excluded because 86% of trials were rejected (see Kinematic Analysis); one participant was excluded as a statistical outlier (see Statistical Analysis).

### Kinematic Analysis

The kinematic data were analyzed in MATLAB (R2021a; The MathWorks Inc., Natick, MA). Movement speed was computed by numerical differentiation of the three-dimensional data divided by the sampling interval of 4 ms. Movement onset was defined as the key release. Mean reaction time was 547 ± 240 ms (standard deviation across participants). Movement offset, i.e., when participants touched the target finger, was determined using the multiple sources of information method to form a composite score (Schot et al., 2010): Coarse spatial constraints excluded frames with lateral positions near the start zone (all positions within 20 cm to the left and 10 cm to the right of the start position were excluded). To lower the likelihood of frames after touching the target finger, frames were down-weighted when the moving index finger was within 3 cm of the location on the table where participants rested their hand after the touch. A timing gate increased the likelihood of a frame being the end of the movement once speed fell below 0.10 m/s after the speed maximum (identified within 400 ms following movement onset) and maintained this elevated score for 80 ms. Lower instantaneous speed, smaller x (toward the target hand), and larger y (further from the body) also increased the score. The movement end was defined as the frame with the highest composite score likelihood.

Movement duration (M = 562 ms, SD = 100 ms) was defined as the time between movement onset and offset. Time of maximum movement speed was determined for each trial. To account for inter-individual differences in movement duration, the timing of maximum speed was normalized to the total movement duration in each trial. Across participants, the average maximum speed was 1.22 ± 0.19 m/s, occurred after 202 ± 62 ms and at 33 ± 6% of the movement. Because maximum speed denotes the onset of the period when sensory guidance of the hand toward the target gains importance (Elliott et al., 2001), stimulation trials were assigned to four movement phases defined relative to maximum speed: (1) pre-max speed (2) window of ± 5% around max speed (3) early post-max speed, the first 40% of the remaining post-max duration; and (4) late post-max speed, the final 60% of the remaining post-max duration. These time bins will be referred to as the four movement phases. We used a 40/60 rather than 50/50 proportion to divide the post-max time to balance trial counts: The average movement duration was 562 ms, while the latest stimulation occurred 450 ms after movement onset, which resulted in too few late-phase trials for stable psychometric fits under a 50/50 post max speed split in eight participants. The 40/60 proportion allowed for stable psychometric fits in seven of these eight participants. Critically, the 40/60 split also yielded more balanced trial counts per movement phase, with phases 1–4 containing on average 32.9%, 13.7%, 27.1%, and 26.3% of trials, respectively. We allowed fewer trials in the small window around maximum speed to maintain temporal specificity in this movement phase.

Trials were excluded from all further analyses if the movement end could not be determined (empty composite score; < 1% of trials), if >25% of motion-tracking samples were missing in the last third of the movement due to marker occlusion (1% of trials), if the mean speed during the first 20% of the movement was below 10 m/s (1% of trials), indicating delayed movement initiation after start key release and incorrectly timed tactile stimulations, if the movement duration was >3 SD above the participant’s within-block mean (1% of trials), or if the stimulation occurred after the movement had ended (4% of trials).

### Psychophysics Analysis

Logistic psychometric functions were fitted to participants’ responses in the psychophysical task using the maximum likelihood estimation implemented in the Psignifit 4 toolbox (Schütt et al., 2016) in MATLAB. Separate functions were fitted for each participant, both for the four movement phases and for the resting condition. Across participants, the mean number of trials included in the rest condition, pre-max, around max, early post-max, and late post-max phases was 70, 108, 45, 88, and 83, respectively. PSEs were derived from the 50% point of each function. To control for inter-individual differences in tactile intensity discrimination, the PSE in each of the four movement phases was subtracted from the corresponding values in the rest condition to compute four normalized difference scores (PSE_diff_) per participant (see Fig. 2).

**Fig. 2:**
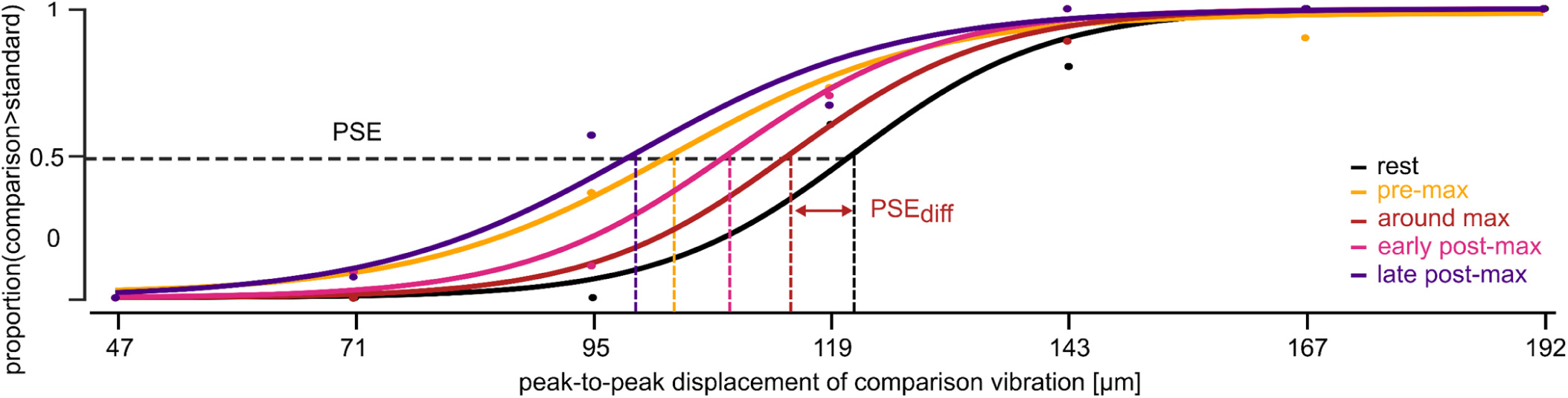
Psychometric curves of one example participant. Psychometric curves were fitted to the participant’s responses in the comparison task, both for the rest condition and for each movement phase. The PSE was determined as the 50% point of each curve. To control for individual differences in tactile sensitivity, the PSE from each movement phase was subtracted from the PSE in the rest condition, yielding one difference score per movement phase (PSE_diff_).

### EEG Analysis

EEG data were recorded from 37 channels (Fp1, C3, P3, P4, C4, AF7, AF3, AFz, F1, F5, FT7, FC3, C1, C5, TP7, CP3, P1, P5, PO7, PO3, POz, PO4, PO8, P6, P2, CPz, CP4, TP8, C6, C2, FC4, FT8, F6, AF8, AF4, F2, FCz) placed according to the international 10–20 system, using a BrainProducts actiCap with a 64-channel LiveAmp system and BrainVision Recorder software (Brain Products GmbH, Gilching, Germany), sampled at 500 Hz. In addition, electrodes were placed at the outer canthi of both eyes and below the left eye to record horizontal and vertical eye movements. Data were processed using eeglab 2024.2 (Delorme & Makeig, 2004) in MATLAB. Preprocessing involved high-pass filtering (non-causal Butterworth impulse response, half-amplitude cut-off at 0.1 Hz, 12 dB/octave roll-off), removal of DC offsets, and attenuation of 50 Hz line noise using Zapline-plus (Klug & Kloosterman, 2022). Data were re-referenced offline to the average of the linked earlobes. Noisy channels were identified through visual inspection and interpolated.

Independent component analysis (ICA) was performed using the Infomax algorithm, with PCA rank reduction used when channels had been interpolated. Ocular artifacts were removed by ICA and ICLabel classification (Pion-Tonachini et al., 2019). Remaining artifacts were rejected via artifact subspace reconstruction. Stimulation triggers were latency-corrected by 4 ms as determined in a dedicated test run of the experiment using a StimTrak system (BrainProducts, Germany) with an audio adapter and a contact microphone attached to the stimulator’s skin contactor. Data were low-pass filtered using a non-causal Butterworth filter (40 Hz half-amplitude cut-off, 12 dB/octave roll-off). EEG data were segmented into epochs from −460 to 400 ms relative to the vibration onset at the possible five stimulation time points (50, 150, 250, 350, and 450 ms post movement onset) and baseline-corrected using the 200 ms pre-stimulus interval. For omission trials, epochs were extracted in the same manner around the five time points (50, 150, 250, 350, and 450 ms post movement onset), to obtain the EEG activity associated with movement at the corresponding vibration delivery times.

These omission epochs were averaged separately for each time point and subsequently subtracted from the respective stimulation epochs, isolating the SEPs (see Fig. 3A). The resulting difference waves were grouped into four movement phases by stimulation timing relative to each trial’s maximum speed (pre-max, around max, early post-max, late post-max; see Kinematic Analysis) and averaged within each phase (see Fig. 3B). Mean trials per participant were 64, 96, 41, 80, and 78 in the rest condition, pre-max speed, around max speed, early post-max speed, and late post-max speed phases, respectively. On average, 61 omission trials were analyzed per person and time point. The analysis focused on electrode CP3 over S1 contralateral to the stimulated right hand. Components were identified individually from each participant’s within-subject average trace across the rest condition and the four movement phases (see Fig. S1). Within predefined ranges, positive peaks were located for the P45 (35–55 ms) and P300 (250–350 ms). Measurement windows were set at ± 5 ms of peak latency for P45 and ± 10 ms for P300. Mean amplitudes were then extracted per participant, rest condition and movement phase within these windows. Next, we computed differences scores by subtracting the mean component amplitude of each of the four movement phases from the mean component amplitude of the rest condition, resulting in one P45_diff_ and P300_diff_ score per movement phase. This rest-movement subtraction was chosen to maintain consistency with the computation of the psychophysical measure.

**Fig. 3:**
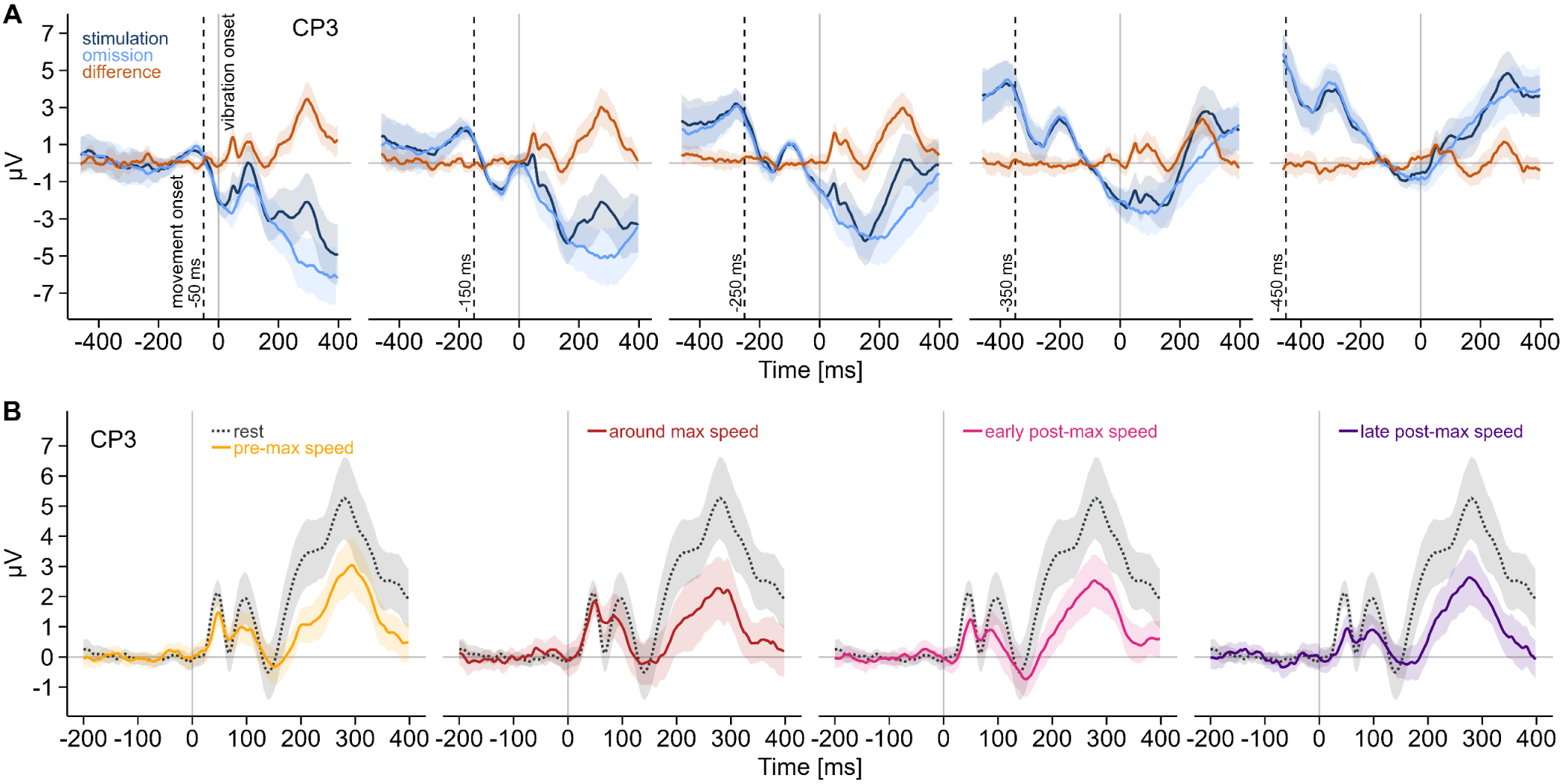
Grand-averaged EEG activity and difference waves across movement phases. **A**. Grand-averaged EEG responses (*n* = 34) for stimulation and omission trials. Stimulation trials with movement and vibrations delivered at five time points (50, 150, 250, 350, 450 ms) post movement onset. EEG was epoched to vibration onset and baseline corrected to the 200 ms pre-stimulation interval. Omission trials with movement only, epoched at the same five time points (50, 150, 250, 350, 450 ms) post movement onset and baseline corrected to the preceding 200 ms. Difference waves computed as stimulation minus omission to isolate SEPs. The dotted vertical line shows movement onset. Omission traces in all subpanels are the same data re-epoched; stimulation traces differ by stimulation time. Shaded bands show 95% CIs. **B**. Grand-averaged difference-wave SEPs (*n* = 34) grouped into four phases relative to maximum movement speed: pre-max speed (yellow), around max speed (red), early post-max speed (pink), late post-max speed (purple). Rest condition SEPs, shown for reference (black dotted), are the same in each subpanel. Shaded bands show 95% CIs.

### Statistical Analysis

To assess whether movement suppressed vibration perception, we compared the PSE_diff_ from each of the four movement phases with zero using paired, two-sided *t*-tests. Modulation of tactile perception across the movement time course was tested with a univariate repeated-measures ANOVA on the PSE_diff_ from the four movement phases. Post hoc comparisons used paired-samples t-tests between movement phases.

To examine modulations in the SEPs associated with movement, we tested P45_diff_ and P300_diff_ from each movement phase against zero using paired, two-sided t-tests. To assess a modulation of component amplitudes across the four movement phases, univariate repeated-measures ANOVAs were performed on P45_diff_ and P300_diff_ separately, followed by paired post-hoc t-tests between movement phases. Planned contrasts that were motivated by significant differences in PSE_diff_ were one-sided; all other comparisons were two-sided. The covariation of perception with changes in SEPs was tested using Pearson correlation. To assess whether SEP amplitudes varied with changes in tactile perception, Pearson correlations were computed between SEP differences and corresponding PSE_diff_ values across phases.

The false discovery rate was controlled within each test family using the Benjamini-Hochberg method, and adjusted *p*-values are reported (Benjamini & Hochberg, 1995). Effect sizes are reported as Cohen’s d_z_ and partial η² (Lakens, 2013). Data were analyzed in R, v.4.3.3 (R Core Team, 2024) with a significance level of α = 0.05. Greenhouse-Geisser correction was applied when sphericity was violated (Mauchly’s test). Univariate outliers were defined as ∣z∣ >3 based on means and standard deviation within movement phases. Bivariate outliers were defined by Mahalanobis distance (*D*^2^) with a cutoff of χ^2^(2, 0.999) = 13.82 (Finch, 2012; Hardin & Rocke, 2005). One participant was classified as both a univariate outlier (*z* = −3.63) within P45_diff_ in the around max speed phase and as a bivariate outlier for P45_diff_ and PSE_diff_ (*D*^2^ = 23.09). Data from this participant were excluded from the primary analyses (see Supplement and Fig. S2 for the analysis without this outlier exclusion).

## Results

To account for individual differences in tactile processing, movement-related changes were expressed as difference scores, obtained by subtracting PSEs and P45 amplitudes obtained during movement from those measured at rest (PSE_diff_ and P45_diff_; see Methods). Positive PSE_diff_ and P45_diff_ values reflect reduced tactile sensitivity and smaller SEP amplitudes, respectively, during movement relative to rest.

To test tactile suppression during movement, we compared PSE_diff_ from each movement phase to zero using paired t-tests. Tactile sensitivity was suppressed relative to rest, with PSE_diff_ significantly greater than zero in all phases of the movement: pre-max speed (*t*(33) = 3.47, *p* = 0.002, *d_z_* = 0.59, 95% CI [5.54–21.24]), around max speed (*t*(33) = 2.48, *p* = 0.019, *d_z_* = 0.42, 95% CI [1.73–17.59]), early post-max speed (*t*(33) = 3.59, *p* = 0.002, *d_z_* = 0.62, CI = [6.36–23.01]), and late post-max speed (*t*(33) = 4.20, *p* < 0.001, *d_z_* = 0.72, 95% CI [9.23–26.60]). A repeated-measures ANOVA used to assess the temporal dynamics of suppression revealed a main effect of movement phase (*F*(3, 99) = 4.66, *p* = 0.004, η²_p_ = 0.12). Post-hoc comparisons showed weaker suppression (i.e., improved sensitivity) around max speed than in the late post-max phase (*t*(33) = −3.3, *p* = 0.014, *d_z_* = −0.57, 95% CI [-13.35– −3.16]), in line with previous findings(Voudouris & Fiehler, 2021). Suppression did not differ significantly between the other movement phases (all *t*(33) ≥ −2.5, *p* ≥ 0.053, *d_z_* ≥ − 0.43; see Fig. 4A).

**Fig. 4:**
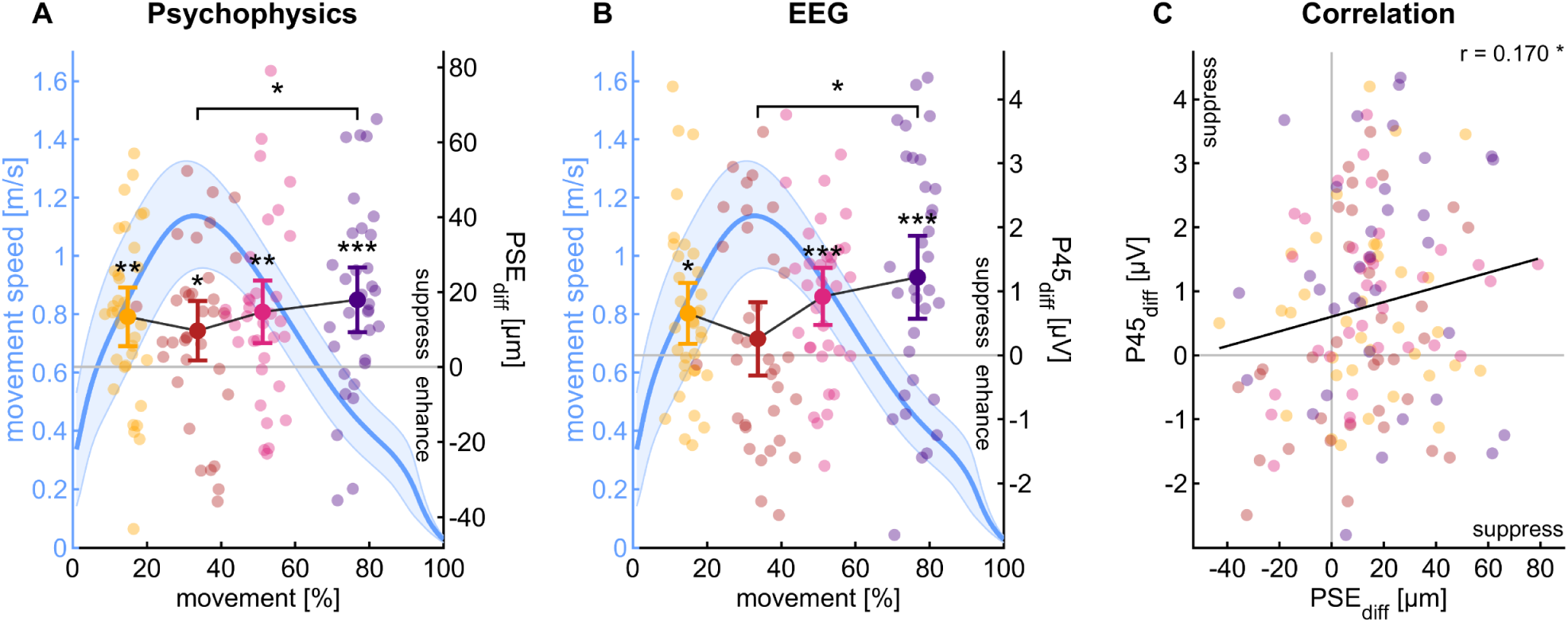
Movement-related suppression in tactile perception and SEP amplitude. Average time-normalized movement speed across participants (*n* = 34; blue line, left *y*-axis). Shaded area indicates standard error. The right *y*-axis shows mean PSE_diff_ (A) and P45_diff_ (B) for the four movement phases: pre-max speed (yellow), around max speed (red), early post-max speed (pink), and late post-max speed (purple). Error bars represent 95% confidence intervals. Dots illustrate individual participants. **A**. Tactile perception was suppressed in all four movement phases (paired *t*-tests against zero). Tactile suppression was weaker around max speed relative to the late post-max speed phase (univariate ANOVA, paired t-tests). **B**. P45 amplitude was comparable to rest around max speed and suppressed in all other movement phases (paired *t*-test against zero). P45_diff_ was smaller around max speed than in the late post-max speed phase, indicating less SEP gating mid-reach relative to later in the movement (univariate ANOVA, paired *t*-tests). **C**. Scatterplot of P45_diff_ versus PSE_diff_. Each point represents one participant in one movement phase. The black line shows the pooled ordinary least squares regression of P45_diff_ on PSE_diff_ across all observations. Stronger tactile suppression during movement was significantly associated with lower P45 amplitude (Pearson correlation). * = *p* < 0.05, ** = *p* < 0.01, *** = *p* < 0.001.

We focused the EEG analysis on electrode CP3 located over S1 contralateral to the stimulated right hand, since P45 amplitude is maximal at this site (Yamada, 2014). Figure 2A shows the waveforms, and Figure 2B the respective scalp maps. To assess SEP gating during movement, we tested P45_diff_ against zero with paired t-tests. P45_diff_ was greater than zero for most movement phases, reflecting gating of the P45: pre-max speed (*t*(33) = 2.78, *p* = 0.010, *d_z_* = 0.48, 95% CI [0.18–1.13]), early post-max speed (*t*(33) = 4.18, *p* < 0.001, *d_z_* = 0.72, 95% CI [0.47–1.36]), and late post-max speed (*t*(33) = 3.83, *p* < 0.001, *d_z_* = 0.66, 95% CI [0.57–1.87]). Importantly, P45_diff_ was not significantly different from zero around maximum speed (*t*(33) = 0.91, *p* = 0.367, *d_z_* = 0.16, 95% CI [-0.32–0.83]), indicating a transient recovery of early cortical processing around the onset of reach deceleration. These findings demonstrate gating of somatosensory signals during movement, except during the time when sensory guidance of the hand gains importance.

A repeated-measures ANOVA revealed a temporal modulation of P45_diff_ across movement phases (*F*(3, 99) = 3.35, *p* = 0.044, η²_p_ = 0.092). Post-hoc t-tests showed significantly smaller P45_diff_ around max speed than in the late post-max phase (*t*(33) = −2.53, *p* = 0.049, *d_z_* = −0.43, 95% CI [-1.73– − 0.19]), supporting the recovery of processing around the time of maximum speed, consistent with the psychophysical results. No significant differences were observed in P45_diff_ between the other movement phases (all *t*(33) ≥ −2.41, *p* ≥ 0.066, *d_z_* ≥ −0.41; see Fig. 4B). Lastly, a significant Pearson correlation between PSE_diff_ and P45_diff_ (*t*(134) = 2.00, *p* = 0.048, *r* = 0.170, 95% CI [0.002–0.33]) revealed that stronger tactile suppression in the PSEs was associated with reduced P45 amplitudes (see Fig. 4C).

Taken together, these findings suggest that early cortical processing contributes to the temporal modulation observed in tactile suppression during reaching. Perceptual reports, however, are shaped not only by early cortical processes, but also by higher-order cognitive mechanisms (e.g., stimulus evaluation and decision-making processes (Heekeren et al., 2008).

To test whether later cortical mechanisms likewise exhibited dynamic changes across the movement, we also examined SEP gating in the P300 component (see Fig. 5). P300_diff_ was significantly greater than zero in all phases, reflecting consistent gating of the P300 throughout the movement: pre-max (*t*(33) = 3.97, *p* < 0.001, *d_z_* = 0.68, 95% CI [1.03–3.21]), around max (*t*(33) = 3.95, *p* < 0.001, *d_z_* = 0.68, 95% CI [1.22–3.81]), early post-max (*t*(33) = 4.81, *p* < 0.001, *d_z_* = 0.82, 95% CI [1.51–3.72]), and late post-max (*t*(33) = 2.85, *p* = 0.010, *d_z_* = 0.49, 95% CI [0.68–4.04]). Additionally, in contrast to the PSE_diff_ and P45_diff_, the P300_diff_ did not significantly differ between the four movement phases (*F*(2.08, 68.59) = 0.43, *p* = 0.658, η²_p_ = 0.013), i.e., we found no temporal modulation at this later stage of somatosensory processing.

**Fig. 5:**
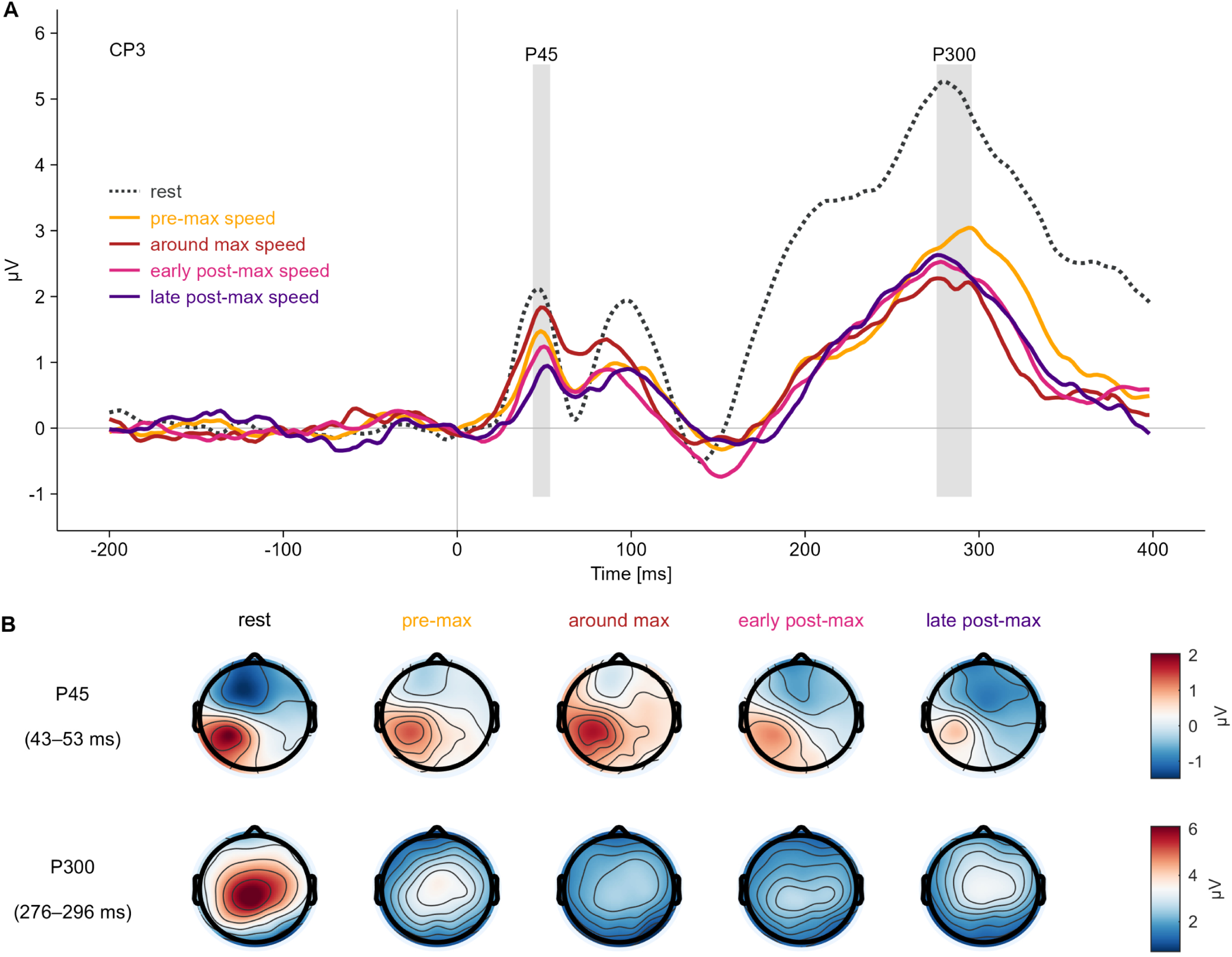
Grand-averaged SEPs during rest and movement. **A**. Grand-averaged SEPs at CP3 (*n* = 34), time-locked to standard vibration onset, with P45 and P300 highlighted. **B**. Scalp topographies for the P45 and P300 across the rest condition and movement phases.

## Discussion

We report temporal dynamics in tactile suppression during goal-directed reaching, with weaker suppression around maximum speed. This temporal modulation was paralleled in early somatosensory processing. As expected, P45 amplitudes were reduced during most movement phases relative to rest, reflecting early cortical gating (Rossini et al., 1999; Rushton et al., 1981). However, P45 responses near maximum speed were comparable to rest and larger than late in the movement. These reductions in P45 amplitude correlated with stronger tactile suppression, suggesting a link between changes in tactile sensitivity during movement and early somatosensory processes. In contrast, P300 amplitudes were uniformly reduced during movement. This suggests that the temporal modulation reflects sensorimotor rather than cognitive processes. Our findings are consistent with reports of phase-dependent sensory modulation during goal-directed movements (Česonis & Franklin, 2020; Dimitriou et al., 2013; Wachsmann et al., 2025a). Similarly, a previous study using the same task reported a complete release from suppression in this movement phase, with tactile sensitivity around maximum speed matching sensitivity at rest (Voudouris & Fiehler, 2021). In contrast, we observed tactile suppression across all movement phases, including maximum speed. This discrepancy may reflect differences in psychophysical task demands: earlier work used a binary detection response immediately after movement offset (Voudouris & Fiehler, 2021), whereas our delayed comparison task required stimulus retention until several seconds after movement completion. Consistent with this, the P300, which is sensitive to memory load (Polich, 2007), was uniformly gated across all movement phases, suggesting that cognitive load may have sustained perceptual suppression, even at phases of preserved early processing. However, the P45 component, a temporally precise index of sensory encoding tied to low-level stimulus features such as intensity (Fuehrer et al., 2025; Schröder et al., 2021), was preserved around maximum speed. Thus, despite stronger overall perceptual suppression in our task, both studies converge on a transient release from suppression around maximum speed. These findings provide a crucial link between perceptual modulation during movement and changes in early sensory processing.

Our results align with optimal feedback control accounts, which propose a state-dependent modulation of feedback gains during movement (Liu & Todorov, 2007; Todorov & Jordan, 2002). Consistent with this, visuomotor gains peak mid-reach and decrease upon approaching the target (Dimitriou et al., 2013). Tactile signals contribute to the transport phase of reaching (Gentilucci et al., 1997), possibly by providing proprioceptive guidance (Aimonetti et al., 2007; Moscatelli et al., 2019). Thus, reduced tactile suppression around maximum speed may reflect a transient upregulation of somatosensory feedback during the transition from movement initiation to feedback-driven deceleration (Elliott et al., 2001). Alternatively, the effect could reflect stronger late suppression as the motor system shifts from position control to movement termination or anticipates target contact (Fraser & Fiehler, 2018; Liu & Todorov, 2007). However, mid-reach P45 amplitudes were comparable to rest, but gated in both earlier and later phases, arguing against a purely late-suppression account and supporting selective preservation of early processing when feedback demands are highest.

The mid-reach reduction in tactile suppression contrasts with reports of higher movement speed increasing suppression (Angel & Malenka, 1982; Cybulska-Klosowicz et al., 2011) and reducing cortical responses (Chapman et al., 1988; Rushton et al., 1981), perhaps due to stronger reafferent masking (Laskin & Spencer, 1979). However, those studies involved isolated joint movements with minimal precision demands. In contrast, suppression during complex movements requiring precision appears unaffected by increased reafferent input from stronger forces (Broda et al., 2020). In our somatosensory-guided task without visual feedback, tactile input likely became more relevant just before the deceleration phase. These findings indicate that suppression is not merely a function of movement parameters like speed, but depends on the task relevance of somatosensory input. This aligns with previous evidence of modulations in short-latency SEPs by movement demands in balancing and precision tasks (Knecht et al., 1993; McIlroy et al., 2003; Wasaka et al., 2017). Similar phase-dependent modulation has been reported in locomotion (Altenmüller et al., 1995; Chapin & Woodward, 1982; Duysens et al., 1990; Mouchnino et al., 2015). Our results further support the link between neural changes and perception by demonstrating phase-dependent SEP modulation during somatosensory-guided reaching.

Unlike traditional SEP paradigms using transcutaneous nerve stimulation (Cruccu et al., 2008), we applied vibrations in the preferred range of Pacinian mechanoreceptors and rapidly adapting type II (RAII) fibers (Bolanowski et al., 1988). This allowed the examination of early cortical responses to naturalistic, low-threshold mechanoreceptor signals, rather than synchronized mixed-fiber activation (Gardner et al., 1984). While the stimulation primarily engaged Pacinian mechanoreceptors, reaching likely generated input from a range of cutaneous and proprioceptive receptors. The P45 component arises mainly from S1 (Allison et al., 1992), which has been associated with early cutaneous-proprioceptive integration (Kim et al., 2015). The observed modulation may therefore reflect broader regulation of somatosensory input, beyond RAII-specific tuning.

Since movement-related gating of somatosensory responses has been documented at multiple stages of the ascending pathway, from first-order synapses in the spinal dorsal column to thalamic nuclei (Chapman et al., 1988; Ghez & Pisa, 1972; Seki & Fetz, 2012), the reductions in P45 amplitude observed here may reflect decreased afferent inflow into S1. At the spinal level, gating is mediated by presynaptic inhibition via local interneurons (Bui et al., 2015). In primates, this presynaptic inhibition is partly driven by descending pathways from the brainstem and cortex and emerges before EMG onset (Seki et al., 2003). Consistent with this, in the rodent whisker system, movement-related gating at the first synaptic stage is abolished when S1, S2, and posterior parietal cortex are lesioned, indicating top-down control of sensory gating already at the first synaptic stage (Chakrabarti & Schwarz, 2018). Converging evidence from humans likewise implicates S1, S2, posterior parietal cortex, and supplementary motor area in downregulating somatosensory perception during movement (Arikan et al., 2021; Haggard & Whitford, 2004). Although descending pathways are likely involved in regulating presynaptic inhibition, stronger gating at cortical than spinal levels in primates suggests additional cortico-cortical modulation (Seki & Fetz, 2012). For example, the motor cortex can recruit inhibitory neurons in the rodent auditory cortex to attenuate self-generated sounds (Schneider et al., 2018), illustrating how motor-related signals can engage local inhibitory circuits in the sensory cortex. Together, these findings support cortical control of tactile feedback gains, with descending pathways regulating afferent inflow and intracortical circuits modulating early cortical responses. The transient mid-reach release from P45 gating may therefore reflect reduced presynaptic inhibition along the pathway, decreased intracortical inhibition, or both. Resolving this will require combined cortical and subcortical measures.

These gating mechanisms operate within a bidirectional sensorimotor control loop in which motor commands shape sensory inflow and somatosensory feedback influences motor output (Azim & Seki, 2019). Cutaneous and proprioceptive signals tune reflexes and shape ongoing movements by regulating motoneuron excitability via spinal interneuron circuits (Bui et al., 2015; Panek et al., 2014). Disrupting presynaptic inhibition of proprioceptive afferents impairs goal-directed reaching in rodents (Fink et al., 2014), whereas blocking cutaneous input increases hand path variability in grasping (Gentilucci et al., 1997), indicating that both excessive and absent somatosensory feedback compromise motor control. Together, these findings suggest that sensory gain is dynamically tuned to task demands. The modulation in early cortical gating observed here likely reflects such adjustments of reafferent gain, suppressing redundant somatosensory input while allowing behaviorally relevant feedback to influence motor circuits (Azim & Seki, 2019). Consequently, changes in perceived tactile intensity during movement may represent the behavioral expression of this distributed sensorimotor gating (Azim & Seki, 2019; McComas, 2016).

### Limitations

One limitation concerns potential overlap of SEPs with task-related events such as movement offset and target contact. This overlap would primarily affect the later P300, whereas the earlier P45 likely occurred during ongoing movement. Because these events were not systematically aligned with stimulation timing, any influence should be random and contribute as noise rather than as systematic bias. To mitigate these effects, we subtracted the EEG signal recorded during reaching alone from that recorded during reach with stimulation. The subtracted signal was time-locked to the corresponding stimulation timing post movement onset and therefore included the same unsystematic movement-related activity present during the stimulation epochs. This approach assumes an additive relationship between movement- and stimulation-related signals, although prior work has questioned this assumption (Quinn et al., 2014). Nevertheless, the resulting SEP waveforms matched canonical patterns reported for vibrations, including a fronto-parietal dipole for the P45, and a centro-parietal positivity for the P300 (Alouit et al., 2025; Fuehrer et al., 2025), supporting the validity of the method.

## Conclusion

In sum, tactile suppression is temporally tuned, with a distinct reduction around maximum speed. This modulation was accompanied by changes in early somatosensory processing, suggesting preserved S1 engagement when somatosensory input from the moving limb is most relevant for the sensory guidance of the hand. This pattern suggests a flexible tuning of suppression and gating according to task demands. Higher-order processes appear to contribute to a general dampening of tactile perception but not to its fine-grained temporal tuning. This novel relation of psychophysical modulation to early neural markers underscores that tactile suppression is a flexible, context-sensitive mechanism coupled to sensorimotor demands.

## Acknowledgments

We thank Malaika Grace Alphonsus, Fabian Krause, Athina Nestoropoulou and Rebecca Oliveri for their valuable help in data acquisition.

## Funding

This work was supported by the Deutsche Forschungsgemeinschaft (German Research Foundation, DFG) as part of the Collaborative Research Centre SFB/TRR 135, Project A4, Grant 222641018, and under Germany’s Excellence Strategy (EXC 3066/1 “The Adaptive Mind”, Project No. 533717223).

## Competing interests

Authors declare that they have no competing interests.

## Author contributions

Conceptualization: EF, DV, LKM, KF

Methodology: EF, DV, LKM, KF

Investigation: EF

Visualization: EF

Supervision: DV, LKM, KF

Writing—original draft: EF

Writing—review & editing: EF, DV, LKM, KF

## Notes

### Competing Interest Statement

The authors have declared no competing interest.

### Summary of Updates

Manuscript revised for length and clarity

