## Supplement for "Temporal dynamics of early somatosensory processing in goal-directed actions"

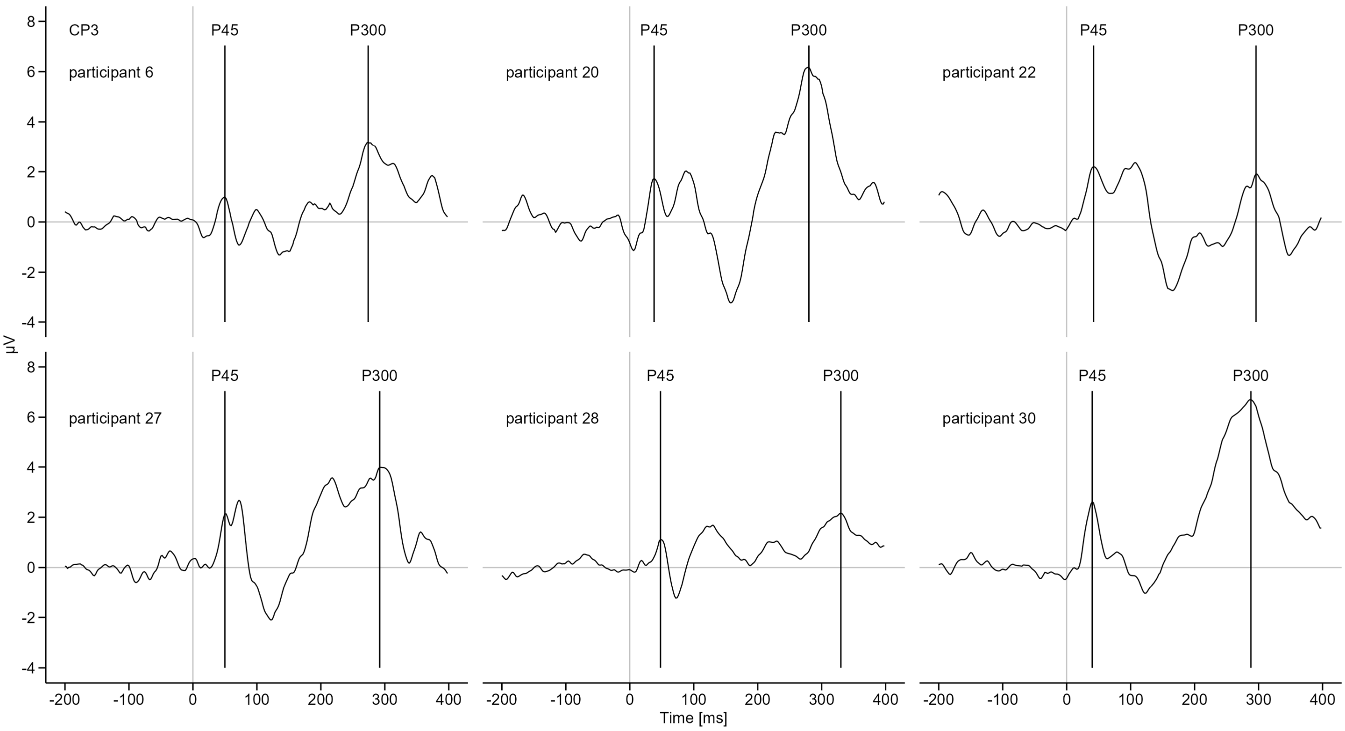


**Fig. S1: Component peak detection.**

Individual waveforms averaged across rest and movement conditions at electrode CP3 for six example participants. Component peaks were identified as maximum values within pre-defined time windows for the P45 (35–55 ms) and P300 (250–350 ms).

**Outlier correction**

One participant was removed after statistical outlier detection. To assess the robustness of the findings, all analyses were repeated with this participant included. All comparisons and main effects remained statistically significant (see Figs. S2A, S2B). One additional post-hoc comparison reached significance: $P45_{diff}$ around max speed differed significantly from $P45_{diff}$ early post-max speed ($t(34) = -2.64$, $p = 0.037$, $d_z = -0.45$; Fig. S2B), indicating higher amplitude of the P45 around maximum movement speed and supporting the main conclusion of stronger suppression later in the movement. In contrast, the correlation between $P45_{diff}$ and $PSE_{diff}$ was sensitive to the inclusion of this participant. With the bivariate outlier included, the correlation did not reach significance $t(138) = 1.09$, $p = 0.277$, $r = 0.092$; see Fig. S2C).

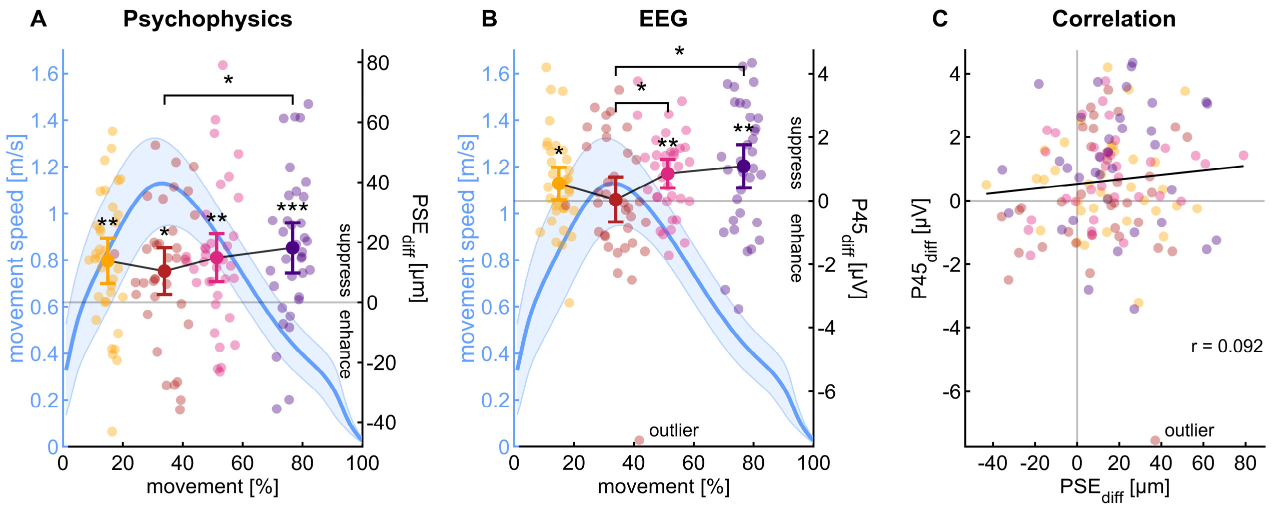


**Fig. S2: Analysis including participant identified as statistical outlier.**

Results are shown for all 35 participants, including the observation of the 1 participant previously excluded based on outlier criteria. The overall pattern of results is preserved. **A**. Tactile suppression ($PSE_{diff}$) was present in all movement conditions (paired *t*-tests vs. zero) and was weaker around maximum speed compared to the late post-max speed phase (paired t-tests). **B**. $P45_{diff}$ was comparable to rest around maximum speed and suppressed in all other conditions (paired *t*-tests vs. zero). In contrast to the main analysis, P45 amplitude was significantly higher around maximum speed compared to both the early and late post-max speed phases. **C**. Scatterplot of $P45_{diff}$ versus $PSE_{diff}$. Each point represents one participant in one movement phase. The black line shows the pooled ordinary least squares regression of $P45_{diff}$ on $PSE_{diff}$ across all observations The previously significant positive correlation no longer reached significance due to the influence of the bivariate outlier (Pearson correlation). * = $p < 0.05$, ** = $p < 0.01$, *** = $p < 0.001$.
